# Controlling Genome Topology with Sequences that Trigger Post-replication Gap Formation During Replisome Passage: The *E. coli* RRS Elements

**DOI:** 10.1101/2023.10.01.560376

**Authors:** Phuong Pham, Elizabeth A. Wood, Emma L. Dunbar, Michael M. Cox, Myron F. Goodman

## Abstract

We report that the *Escherichia coli* chromosome includes novel GC-rich genomic structural elements that trigger formation of post-replication gaps upon replisome passage. The two nearly perfect 222 bp repeats, designated Replication Risk Sequences or RRS, are each 650 kb from the terminus sequence *dif* and flank the Ter macrodomain. RRS sequence and positioning is highly conserved in enterobacteria. At least one RRS appears to be essential unless a 200 kb region encompassing one of them is amplified. The RRS contain a G-quadruplex on the lagging strand which impedes DNA polymerase extension producing lagging strand ssDNA gaps, ≤2000 bp long, upon replisome passage. Deletion of both RRS elements has substantial effects on global genome structure and topology. We hypothesize that RRS elements serve as topological relief valves during chromosome replication and segregation. There have been no screens for genomic sequences that trigger transient gap formation. Functional analogs of RRS could be widespread, possibly including some enigmatic G-quadruplexes in eukaryotes.

## INTRODUCTION

The *E. coli* chromosome is maintained as a highly condensed nucleoprotein structure called a nucleoid. Condensation of the nucleoid arises from several factors (reviewed in (1)). These include nucleoid-associated proteins and structural maintenance of chromosome proteins that bind and condense DNA at both local and global levels (2). Additionally, DNA supercoiling arising from processes such as RNA transcription contribute to nucleoid condensation (3).

Nucleoid condensation in *E. coli* is neither random nor uniform. The *E. coli* chromosome is divided into 31 self-interacting segments called chromosome interaction domains or CIDs (4–10). At least some of the CID boundaries correspond to highly transcribed genes. In addition to CIDs, the genome is further structured into several macrodomains with boundaries that prevent the diffusion of supercoiling and restrict interaction between macrodomains (6–10). Nothing is known about the nature of the structure that defines macrodomain boundaries. One macrodomain (TER) encompasses a million or more bp surrounding the replication terminus defined by the sequence *dif* (11,12).

In addition to genes that encode functional proteins and RNA molecules, the *E. coli* genome includes a variety of genomic elements that facilitate some aspect of genome structure and function. These include the replication origin, *ori*C; the replication terminus region with its *ter* and *dif* sites (13); and about 300 bacterial interspersed mosaic elements (BIMEs). One family of BIMEs functions as efficient binding sites for DNA gyrase (14).

There have been multiple efforts to screen the *E. coli* genome for repeated sequences of many types (15–19). Most of the targeted repeats were transient gene duplications that are unstable and ultimately eliminated by recombination, inverted and tandem intergenic repeats that often have some role in gene expression, transposons, simple sequence repeats, genes with multiple genomic copies, and CRISPR repeats. We report here on a repeated genomic sequence that does not fit into any of these categories.

We recently developed a new method to examine protein binding to single-stranded DNA on a genomic scale, designated ssGAP-seq (20). In applying it to RecA and SSB binding to ssDNA, we identified two sites in the *E. coli* genome where both proteins were deposited at elevated rates on the lagging strand template. At the ends of both sites, we identified a pair of 222 bp and GC-rich repeats. These are positioned symmetrically about the terminal *dif* site, each 650 kb distant from *dif*. When cloned, the repeats cannot be sequenced by standard commercial sequencing unless protocols with extra heat are applied. In the genome, the repeats are causing post-replication gap formation as replisomes pass. We have therefore named them Replication Risk Sequences or RRS (20). The RRS appeared briefly in a 1999 screen for genomic repeats (15) and designated TRIP1 (for TRIPle; a third partial repeat was identified to make 3). However, no functional significance was attached to the repeats and they have been otherwise overlooked.

The RRS are highly conserved in both positioning and sequence among the enterobacteria (20), although some species have one or more additional RRS within their genomes. The conservation and effects on replication suggest a sequence of genomic functional significance. The present study introduces important properties and genomic effects of the RRS elements.

## MATERIALS AND METHODS

### Materials

*E. coli* MG1655, CSH50, MC1061, isogenic RRS deletion strains, RRS inverted strains, mutant strains containing extra RRS sequences, bacteriophage M13mp2, purified wild-type *E. coli* DNA polymerases II (21) and IV (22) and polyclonal chicken anti-RecA and anti-SSB antibodies were from our labs. PrecipHen (Agarose coupled Goat anti-Chicken IgY beads) was purchased from Aves Labs, inc (Davis, CA). EZ-Link Psoralen-PEG3-Biotin and Dynabeads MyOne Streptavidin T1 were from ThermoFisher. EcoRI, MfeI restriction enzymes, T4 DNA ligase, Klenow fragment (3’→5’ exo^−^) and Monarch PCR & DNA clean-up kit was purchased from New England Biolabs. DNase-free RNAse A (100 mg/ml) was from Qiagen, Proteinase K, sodium metabisulfite and hydroquinone were from Sigma-Aldrich. xGen Methyl-Seq Library kit was purchased from Integrated DNA Technologies and Nextera XT library prep kit was from Illumina, Inc.

### Construction of double rΔRRS deletion strain EAW1710

The region from 168bp-389bp upstream of the *lysO* gene = the *lysO* RRS was deleted from MG1655 using λRED recombination. The deletion was confirmed by PCR using primers outside the RRS and direct sequencing of the PCR product. The strain was designated EAW 1706. The region from 281 bp-58 bp downstream of the *dusC* gene = the *dusC* RRS was deleted from MG1655 using λRED recombination. The deletion was confirmed by PCR using primers outside the RRS and direct sequencing of the PCR product. The strain was designated EAW1708. The KanR gene replacing the *lysO* RRS of EAW1706 was removed by FLP recombination, and P1 transduction was used to add the *dusC* RRS deletion from EAW1708. Two colonies from the transduction had different morphologies and were designated EAW1710 colony #1 and #3. Both *Δ*RRS deletions were confirmed by PCR and whole genome sequencing.

### Construction of strains with extra RRS elements

The region from 168 bp-434 bp upstream of the *lysO* gene = extended *lysO* RRS was cloned into a plasmid next to the FRT-KanR-FRT cassette used in λRED recombination. The resulting plasmid, designated pEAW1330, was used as template in PCRs with primers designed to insert the extended *lysO* RRS+ FRT-KanR-FRT cassette 129 bp upstream of the *fabR* gene start codon, 6 bp downstream of the *djlC* gene stop codon, and 72 bp downstream of the dinI gene stop codon. P1 transduction was used to add the RRS near *djlC* to the KanS strain with an extra RRS near *fabR*, and the KanR was removed by FLP recombinase. The extra RRS near *dinI*, and deletions of the wild-type RRS near *lysO*, then the wild-type RRS near *dusC* were made by subsequent P1 transductions. The resulting strain is designated EAW1776, and has 3 extra RRS near *fabR*, *djlC*, and *dinI*, with both wild-type RRS deleted. After each P1 transduction, the extra RRS and wild-type RRS deletions were confirmed by PCR using primers outside the insertion or deletion area and direct sequencing of the PCR product.

### Construction of strains with RRS elements in the reverse orientation

The region from 168 bp-434 bp upstream of the *lysO* gene = extended *lysO* RRS was cloned into a plasmid next to the FRT-KanR-FRT cassette used in λRED recombination, in the reverse orientation as the RRS in plasmid pEAW1330. The resulting plasmid was designated pEAW1331 and was used as template in PCRs with the same primers used with pEAW1330. These primers were designed to insert the extended *lysO* RRS+ FRT-KanR-FRT cassette 129 bp upstream of the *fabR* gene start codon, 6 bp downstream of the *djlC* gene stop codon, and 72 bp downstream of the dinI gene stop codon.

### Cloning of RRS elements into M13mp2 phage vector

*RRS-lysO* and *RRS-dusC* sequences were amplified from MG1655 genomic DNA using Q5 High-Fidelity master mix (New England Biolabs) and pairs of primers containing a MfeI restriction site at each end. The PCR products were digested with MfeI to generate RRS fragments with AATT-5’ overhangs that are compatible with EcoRI digested ends. M13mp2 DNA (RF II form) was linearized at a single site in *lacZ*α reporter gene by EcoR1 restriction enzyme and ligated with a MfeI-digested RRS fragment using T4 DNA ligase at 16 °C for 16 h. Ligated DNA was then introduced into *E. coli* MC1061 competent cells and plated on a bacterial lawn of CSH50 host cells in the presence of X-gal and IPTG as described previously (23). White M13 phage plaques representing M13 recombinant phage containing an RRS insert were individually picked, amplified overnight at 37 °C, and phage ssDNA were purified. The presence of RRS-G4 strand (or RRS-C strand) in recombinant ssDNA phage genome was verified by direct Sanger sequencing, or by PCR screening. PCR primer pair RRS-1F (5’-GATGTTACCTCGCGCCCGA-3’) and LacZ6 (5’-CGCCAGCTGGCGAAAGGG-3’) was used for screening of M13-RRS-G4 phage, whereas primer pair RRS-1R (5’-TCGGGCGCGAGGTAACATC-3’) and LacZ6 was used for screening for M13-RRS-C phage.

### Measurement of relative production of wild-type and recombinant M13 phages

A mixture containing 2.5 ng of wild-type M13mp2 and 2.5 ng of a recombinant M13-RRS-G4 (or M13-RRS-C) dsDNA was introduced into *E. coli* MC1061 competent cells by electroporation and plated on a bacterial lawn of CSH50 host in the presence of X-gal and IPTG as described previously (23). After overnight incubation at 37 °C, wild type M13 phages were scored as dark blue plaques and recombinant phages were scored as white (colorless) plaques.

### Polymerase primer extension

M13seq primer (5’-GACGTTGTAAAACGACGGC-3’) was labeled with ^32^P at 5’-end by T4 polynucleotide kinase (New England Biolabs). Primer/template (p/t) substrates were prepared by heating a tube (50 μl of 10 mM Tris-HCl, pH 7.9, 50 mM NaCl) containing 40 nM labeled M13seq primer and 50 nM purified M13, M13-RRS-G4 or M13-RRS-C ssDNA at 99 °C for 2 min and slowly cooled down to room temperature. Primer extensions were carried out at 37 °C for 30 min in a reaction buffer (10 mM Tris-HCl, pH 7.9, 50 mM NaCl, 10 mM MgCl_2_ and 1 mM DTT) containing 2 nM of a p/t substrate, 500 μM dNTP and a polymerase (0.5 units of Klenow fragment exo-, 50 nM of Pol II or 500 nM of Pol IV). After stopping reactions by addition of 20 mM EDTA, extension products were separated by electrophoresis on a 10% denaturing PAGE gel and visualized by a Typhoon Phosphorimager (GE Healthcare, Inc).

### ssGap-seq

Measurement of the genome-wide distributions of RecA and SSB bound to ssDNA by ssGap-seq was carried out essentially as described in Pham et al (20).

### Psora-seq

Measurement of whole-genome supercoiling at high resolution by Psora-seq was carried using a published protocol (24) with several modifications detailed below.

### Cell preparation for Psora-seq

Cells from wild-type MG1655, mutant EAW1706 *(ΔRRS-lysO*), EAW1708 *(ΔRRS-dusC*) and EAW1710 (double deletion *ΔRRS-lysO*, *ΔRRS-dusC*) strains were grown in LB medium at 37 °C. 100 ml of cells in mid-exponential phase (OD_600_=0.2) were chilled on ice for 3 min, then harvested by centrifugation (4000 x g, 5 min) and resuspended in 10 ml of ice-cold TE buffer (10 mM Tris, pH 7.4, 1 mM EDTA) containing 5 μg/ml biotinylated psoralen (EZ-Link Psoralen-PEG3-Biotin, ThermoFisher). Cell suspensions were incubated in the dark at 4 °C for 10 min, then transferred to a 100 mm polystyrene Petri dish. Crosslinking of psoralen to genomic DNA was performed by cell exposure to 365 nm UV light (4.8 J/cm^2^ total energy) using a UVP CL-3000L crosslinker (UVP, Inc. Upland, California). Cells were washed with 40 ml of ice-cold TE buffer and cell pellets were stored at −80 °C.

### Genomic DNA purification

Psoralen-crosslinked cells were lysed with lysozyme (2 mg/ml) in 4 ml of lysis buffer (20 mM Tris, pH 8.0, 5 mM EDTA, 100 mM NaCl) for 10 min at 37 °C. Cellular RNA was removed by treatment with RNAse A (100 μg/ml) for 30 min at 37 °C. Cellular proteins were then digested by addition of Proteinase K (final concentration 0.5 mg/ml) and SDS (final concentration 1%) and incubated for 2 h at 37 °C. Genomic DNA (gDNA) was purified by twice extracting with phenol:chloroform: isoamyl alcohol (25:24:1) and precipitated with ethanol. Purified gDNA were resuspended in 250 μl of TE buffer (10 mM Tris-HCl, pH 8.0; 0.1 mM EDTA) and stored at −20 °C.

### Psoralen affinity purification

Genomic DNA (120 μl volume) was sheared to an average size of 400-500 bp by sonication using a bath sonicator (Covaris S2) in a 6 x 16 mm microTUBE AFA fiber. The temperature of the water bath was held between 5 to 8 °C. Sheared DNA was quantified by Qubit Fluorometer (ThermoFisher). For affinity purification, 50 μl of 1 mm diameter streptavidin-coupled magnetic beads (Invitrogen Dynabeads MyOne Streptavidin T1) were washed 3 times and incubated with 10 μg of sheared gDNA in 1 ml B&W buffer (5 mM Tris, pH 7.5, 0.5 mM EDTA, 1 M NaCl, 0.05% Tween-20) for 1 h at room temperature, followed by 18 h at 4 °C in the dark. Unbound DNA was removed by washing 6 times with B&W buffer and 2 times with TE buffer at room temperature. Washed beads were resuspended in 100 μl of TE, and DNA was eluted by twice extracting with an equal volume of phenol:chloroform:isoamylalcohol (25:24:1). The phenol solution efficiently disrupts biotin-streptavidin interaction, allowing high DNA recovery approaching 100% at room temperature (25). Psoralen pull-down DNA was further purified using a Monarch PCR & DNA clean-up kit (New England Biolabs), eluted in 15 μl of EB buffer (10 mM Tris, pH 8.5) and stored at −20 °C.

### Genome sequencing and analysis

1 ng of psoralen pull-down DNA (or corresponding Input DNA from the same cells without pull-down) was used to prepare sequencing libraries using a Nextera XT library prep kit (Illumina, Inc) according to the manufacturer protocol. Libraries were quantified by Qubit, qPCR and subjected to Illumina’s sequencing (2 x 150 bp paired-end) on a MiniSeq instrument using High Output Reagent Kits (300-cycles).

Illumina sequencing data were analyzed using the CLC Genomic Workbench (v. 23.0) (Qiagen). High quality sequencing reads were imported, 5 nt were trimmed from each end, and mapped to an MG1655 reference genome (Accession number NC_000913) using read alignment settings of 0.8 for a length fraction and 0.9 for similarity fraction. Default settings for match score, mismatch, insertion and deletion costs were used. Mapping coverages of aligned reads (1 - 3 million per sample) with detailed per-base coverage information for psoralen pull-down reads and input reads were exported from the CLC Workbench program in a tab delimited format.

Genome-wide binding of psoralen was calculated as described by Visser et al. (24) as follow. A custom-written R-script was used to calculate the normalized read coverages for the psoralen pull-down and the corresponding Input samples at individual positions by taking moving averages per 1 kb window. Psoralen enrichment (z-scores) was calculated by calculating the ratios of normalized psoralen pull-down values to normalized Input values then log_2_ transformed (24). Psoralen binding profiles on larger scales (5 kb, 10 kb, 25 kb, 50 kb, 100 kb, or 250 kb) were calculated similarly, using normalized read coverage values for a moving window of 5 kb, 10 kb, 25 kb, 50 kb, 100 kb, or 250 kb, respectively. To compare psoralen binding profiles of wild-type and RRS deletion strains, psoralen binding z-scores at genomic positions from independent replicates were averaged, and then used to calculate Pearson correlation coefficients (*r*). R-programming was used for statistical analyses and to generate psoralen binding graphs.

## RESULTS

### RRS elements contain a G-quadruplex forming sequence, which obstructs DNA polymerase progression in vitro and in vivo

The RRS elements are 222 bp in length, positioned in the *E. coli* genome near the *dusC* and *lysO* genes (Figure 1A) (20). These sites are each approximately 650 kb from the *dif* sequence that helps define replication termination. The RRS are GC-rich and have the potential to form extensive secondary structure. Using a G-quadruplex (G4) predictor tool (26), we identified a single putative G4 forming sequence on the sense RRS strand (RRS-G4 strand) with a high confidence cG/cC score (27) of 11.3 (Figure 1B). No G4-forming sequence was identified on the RRS complementary anti-sense strand (RRS-C strand). Notably, due to RRS orientations in the *E. coli* genome, the G4 forming strand of RRS at both *lysO* and *dusC* loci serves as the lagging strand template during chromosomal replication (Figure 1A). An *E. coli* genome search for the G4 sequence (5’-GGGGGTGCAGGGGGCGGCGG-3’) did not yield any additional site.

**Figure 1.**
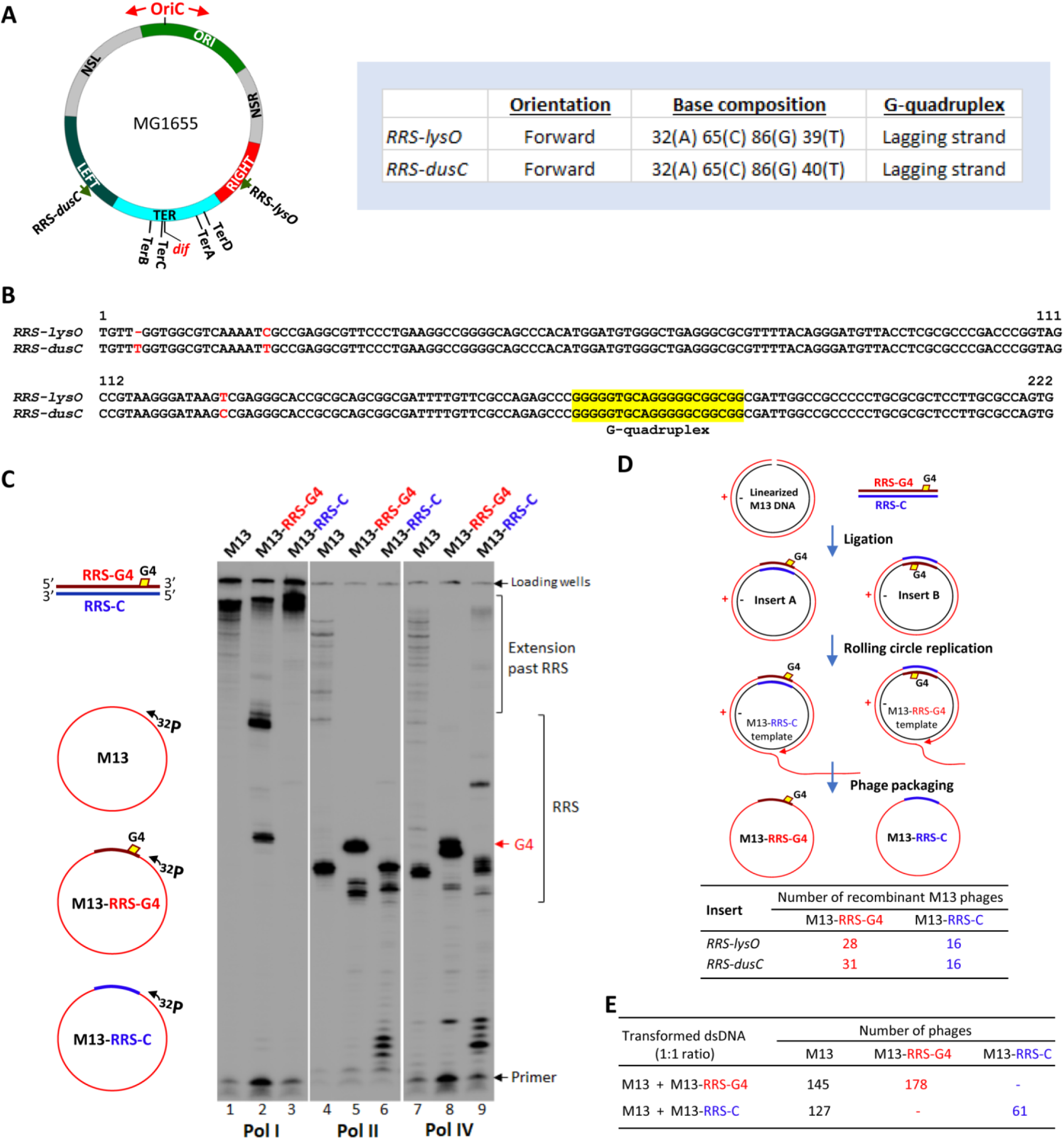
G-quadruplex (G4) on the RRS sense strand blocks DNA polymerase extension. **A**, Locations and base composition of RRS at *lysO* and *dusC* loci. Positions of RRS-*lys*O, RRS-*dusC*, OriC, *dif, Ter* sites, four macrodomains (ORI, LEFT, RIGHT, TER) and two non-structured domains (NSL and NSR) of *E. coli* chromosome are shown. **B**, Alignment of RRS-*lysO* and RRS-*dusC* sequences. A single G4-forming sequence was identified on the RRS sense strand, but not on the anti-sense strand (yellow highlight). **C**, RRS G4 blocks *E. coli* DNA polymerase extension *in vitro*. Extension of a ^32^P labeled primer by Pol I (Klenow exo-), Pol II and Pol IV was performed using three circular phage ssDNA substrates: wild-type M13mp2 (M13), recombinant phage containing the RRS sense strand with G4 (M13-RRS-G4), and phage with the inserted RRS complementary strand (M13-RRS-C) (*sketch on the left*). Strong Pol I, Pol II and Pol IV extension stopping bands are observed at the G4 site on M13-RRS-G4 substrate (lanes 2, 5 and 8, G4 arrow). **D**, Less efficient M13 rolling circle replication by Pol III holoenzyme *in vivo* when RRS G4 is present on M13 (-) template strand. Random ligation produces equal numbers of ligated M13-RRS molecules in forward and reverse orientations (*sketch on top*). Less efficient rolling circle replication for ligated molecules with the G4 present on the (-) template strand resulted in a significantly fewer number of recombinant M13-RRS-C phage produced, compared to M13-RRS-G4 phage (*p < 0.05*, Chi square test). **E**, Production of wild-type M13 and recombinant phages (M13-RRS-G4 or M13-RRS-C) upon transformation of a mixture of wild-type M13 and recombinant phage dsDNA (1:1 ratio) into *E. coli* cells.

To gain additional insight into RRS functions, specifically for the G4 forming sequence, we examined RRS G4 effects on DNA synthesis by four *E. coli* DNA polymerases: Pol I, Pol II, Pol IV *in vitro*, and Pol III holoenzyme *in vivo* (Figures 1C, 1D & 1E). A circular ssDNA genome of M13mp2 phage (M13) along with recombinant phage ssDNA containing either the RRS-G4 strand (M13-RRS-G4) or the complementary RRS-C strand (M13-RRS-C) were annealed to a ^32^P labeled primer and used as primer/template substrates for polymerase extension by Pol I (Klenow Fragment exo-), Pol II and Pol IV (Figure 1C). Pol I exhibits a robust activity on M13 and M13-RRS-C substrates, synthesizing full-length of M13 DNA without a visible pausing band (lanes 1 & 3, Figure 1C). However, on M13-RRS-G4 substrate, a strong Pol I pausing band at the G4 site and upper bands located at the end of RRS-G4 insert are observed (lane 2). Pol II and IV can extend the primer to the end of control M13 ssDNA substrate, but both enzymes pause strongly at positions ∼30 nt downstream from the primer (lanes 4 and 7), suggesting that these polymerases are quite sensitive to a secondary structure in the control M13 substrate (28). DNA synthesis by both Pol II and IV is completely blocked at the G4 site on M13-RRS-G4 substrate (lanes 5 and 8). The RRS-C strand completely blocks DNA synthesis by Pol II at a different location within the RRS, but Pol IV blockage is only partial (lanes 6 & 8).

Thus, while DNA synthesis by Pol I, II and IV is strongly blocked at the G4 site on the RRS-G4 strand, the RRS-C strand appears to have differential and smaller effects on the DNA polymerases.

We further explored the impact of the RRS G4 sequence for DNA synthesis *in vivo* using M13 rolling circle replication carried out by Pol III holoenzyme in *E. coli* (29). We used T4 ligase to ligate a RRS fragment (RRS-*lysO* or RRS-*dusC*) into linearized M13 dsDNA vector. Random ligation results in equal numbers of recombinant M13 molecules containing a RRS insert in the forward orientation (Insert A with the G4 on M13 (+) strand) and the reverse orientation (Insert B with the G4 on M13 (-) strand) (Figure 1D). Upon transformation into *E. coli*, the M13 ssDNA genome is replicated by rolling circle replication using the (-) strand of dsDNA as the template (29). Phage packaging produces M13 recombinant phage containing an RRS-G4 or RRS-C strand. Equal numbers of recombinant phage M13-RRS-G4 and M13-RRS-C would be expected, if the RRS G4 does not have any effect on Pol III replication. We screened 44 recombinant phage clones with RRS-*lysO* and 47 clones with RRS-*dusC* and found that the number of recombinant M13-RRS-G4 phage is nearly 2-fold higher (*p < 0.05*) than the number M13-RRS-C phage (Figure 1D), indicating that fewer phages were produced when the RRS G4 is present on the (-) template strand. Since Pol III performs leading strand synthesis during M13 genome replication, the data suggest that the presence of RRS G4 lead to about 2-fold reduction in Pol III holoenzyme leading strand synthesis in *E. coli*, relative to the RRS-C strand.

Another experiment was carried out that included the wild-type M13mp2 DNA as a control to determine if the RRS-C strand also has a detectable effect on Pol III-mediated leading strand synthesis. We introduced a mixture of phage dsDNA from M13 and a recombinant phage (either M13-RRS-G4 or M13-RRS-C) at 1: 1 ratio into *E. coli* cells and monitored production of wild-type M13 and recombinant phages. Data in Figure 1E showed that while the number of M13-RRS-C phage produced is about 2-fold lower than the number for wild-type, the number of M13-RRS-G4 phage produced is slightly higher than the number for wild-type (Figure 1E), suggesting that the RRS-C strand does not have a detectable effect on Pol III-mediated rolling circle replication.

### RRS with the G4 on the lagging strand triggers ssDNA gap formation wherever they occur

RRS were originally defined as sites that anchored a prominent peak in RecA and SSB deposition on single-stranded DNA (20). The RecA is deposited uniquely on the lagging strand template, in a region of approximately 2 kb (Figure 2B). Deposition is greatest near the RRS element and trails off from that point. This behavior suggests that DNA polymerase subunits replicating the lagging strand are halted at the RRS G4 site, disengage, and re-initiate downstream, leaving a gap into which SSB and/or RecA is often loaded.

**Figure 2.**
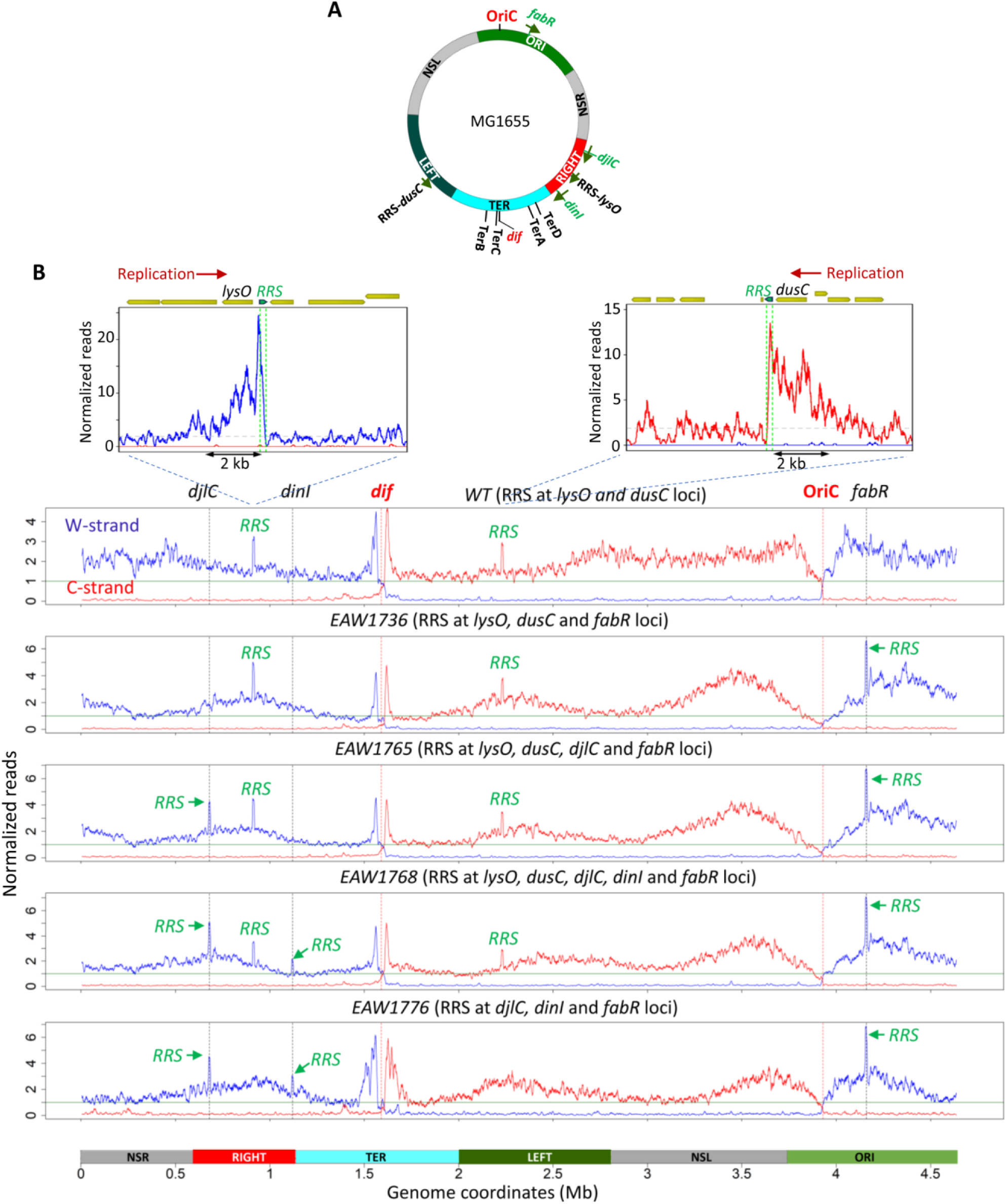
RRS induces formation of genomic ssDNA on the lagging strands. **A.** Sketch depicting positions of RRS-*lys*O, RRS-*dusC*, OriC, *dif, Ter* sites, four macrodomains (ORI, LEFT, RIGHT, TER) and two non-structured domains (NSL and NSR) of *E. coli* chromosome. Positions of *fabR*, *djlC* and *di*nI genes are shown in green color. **B,** Genome-wide distribution of RecA-ssDNA in *wild-type* and mutant strains containing extra RRS sequences inserted at *fabR* locus (EAW1736); *fabR* and *djlC* loci (EAW1765); *fabR, djlC* and *dinI* loci (EAW1768). Strain EAW1776 contains three RRS elements at *fabR, djlC* and *dinI* loci, but two original RRS at *lysO* and *dusC* had been removed. Normalized coverages of RecA-ssDNA are shown as blue traces for the W-strand, and red traces for the C-strand. Green lines in graphs (value = 1) indicate the genome average coverage. Each data point represents a moving average of RecA-ssDNA in a 10 kb window. *Top inserts*, zoom-in RRS-*lysO* and RRS-*dusC* regions showing RecA-ssDNA binding on the lagging strand peaks at RRS then extends further by about 2 kb.

The positions of these sequences, near the reported boundaries of the Ter macrodomain (11,12), suggested an important function. We have pointed out that the RRS sequences are highly conserved in both sequence and positioning among enterobacterial species (30,31), although a few species have additional RRS. We wished to determine if the *E. coli* RRS elements maintained their effects if they were positioned in other parts of the genome. We therefore constructed several strains in which one or both of the RRS were deleted or in which RRS elements were placed in other genomic locations in the same orientation as the natural RRS relative to the direction of replisome travel. A series of strains were constructed in which RRS elements with the RRS G4 on the lagging strand were placed near the *fabR*, *djlC*, and *dinI* genes, singly or together (see locations in Figure 2A). All of these strains, with one, two, or three new RRS, were subjected to ssGAP-seq to determine where RecA and SSB were deposited on ssDNA. In every case, when a new RRS was positioned in the genome, a new peak of RecA and SSB deposition appeared at that location (see results for EAW1736, 1765, and 1768 in Figure 2B and Supplementary Figure S1). In each case, the RRS was oriented the same way so that the RRS G4 strand represents the lagging strand template of a traveling replisome. The results indicate that the RRS G4 will function as lagging strand gap-triggering elements wherever they are positioned in the genome. We note that the presence of additional RRS elements did have significant effects on the global pattern of generation of lagging strand ssDNA and RecA deposition, especially in the 1 Mb or so near the replication origin (Figure 2B). We cannot yet comment on possible molecular events underlying these effects.

### RRS elements appear to function in the reverse orientation, with limits

To examine the effects of RRS element orientation on genome function, we constructed a series of strains in which one or both RRS were inverted. The strains with one inverted RRS were in some cases constructed with the second RRS deleted. In all cases, construction of strains with inverted RRS was possible and the strains grew well. For inverted RRS paired with RRS deletions, the RRS inversion had to be in place before the deletion of the other RRS was attempted. We carried out ssGap-seq for RRS inverted strains (Figure 3 and Supplementary Figure S2). As was the case for adding new RRS in new locations, the inversion of one or both RRS elements had substantial effects on the pattern of ssDNA generation and RecA deposition particularly in the genomic domains near the replication origin (Figure 3 and Supplementary Figure S2).

**Figure 3.**
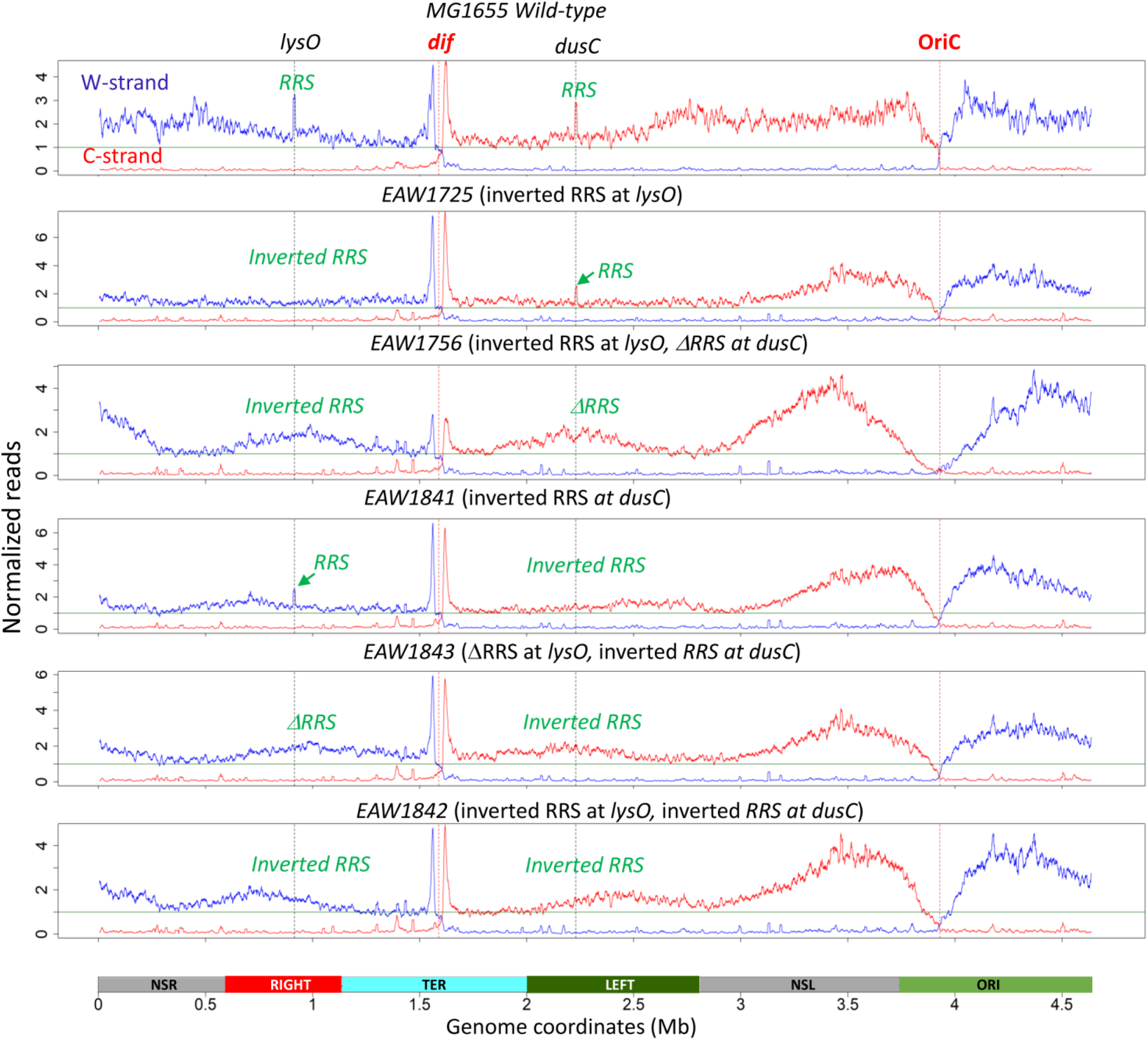
Genome-wide distribution of RecA-ssDNA in *wild-type* and mutant cells with inverted RRS. Mutant strains contain an inverted *RRS-lysO* (EAW1725), an inverted *RRS-lysO* and *DRRS* deletion at *dusC* (EAW1756), an inverted *RRS-dusC* (EAW1841), an inverted *RRS-dusC* and *DRRS* deletion at *lysO (*EAW1843), and double inverted *RRS-lysO* and *RRS-dusC* (EAW1842). Normalized coverages of RecA-ssDNA are shown as blue traces for the W-strand, and red traces for the C-strand. Green lines in graphs (value = 1) indicate the genome average coverage. Each data point represents a moving average of RecA-ssDNA in a 10 kb window.

In contrast to RRS in the forward orientation, neither RecA-ssDNA or SSB-ssDNA peak was observed at the inverted RRS loci (Figure 3 and Supplementary Figure S2), suggesting that RRS in the reverse orientation did not trigger ssDNA gap formation or at least did so less frequently or without RecA loading. RRS inversion re-positioned the RRS-G4 on the leading strand of a moving replication fork. The ability of the RRS-G4 to trigger ssDNA gap formation when present on the lagging strand, but not on the leading strand is likely attributed to the discontinuous nature of lagging strand synthesis. A lagging strand ssDNA gap region is expected to form between the RRS-G4 polymerase blocking site and the 5′-end of the preceding Okazaki’s fragment, whereas the leading strand Pol III probably stalls at the RRS-G4 site and resumes DNA synthesis after G4 structure resolution by SSB or a helicase without generation of ssDNA gaps (32). Additional effects of RRS inversion on genome topology are documented below.

### RRS elements are conditionally essential for genome function

Deletion of one or the other natural RRS does not lead to any measurable cell stress or growth defect. However, we have been unable to construct a clean deletion of both RRS. Multiple attempts to delete the RRS near *dusC* in a strain lacking the *lysO* RRS resulted in a few colonies that grew as well as wild type cells. However, in all cases, genome sequencing revealed that these strains have an amplification of the 200 kb genomic region between *insH7* and *insH8* (Figure 4, Supplementary Figure S3A). This region encompasses the *dusC* RRS element. For all double Δ*RRS* colonies, the average number of sequencing reads in the *insH7-8* region is 2.3- to 3.0-fold higher in the genome average (Figure 4, Supplementary Figure S3A). Analysis of the *insH7-8* genomic region by PCR using primers flanking *insH7* and *insH8* showed that surviving double ΔRRS deletion colonies contain a tandem duplication or triplication of the *insH7-8* region (Supplementary Figure S3B). Whole genome mutation analysis for single and double Δ*RRS* deletion mutants did not identify any mutation that is specific for double Δ*RRS* deletion colonies (Supplementary Figure S4). We conclude that at least one RRS is essential unless its function is somehow compensated for with this genome rearrangement.

**Figure 4.**
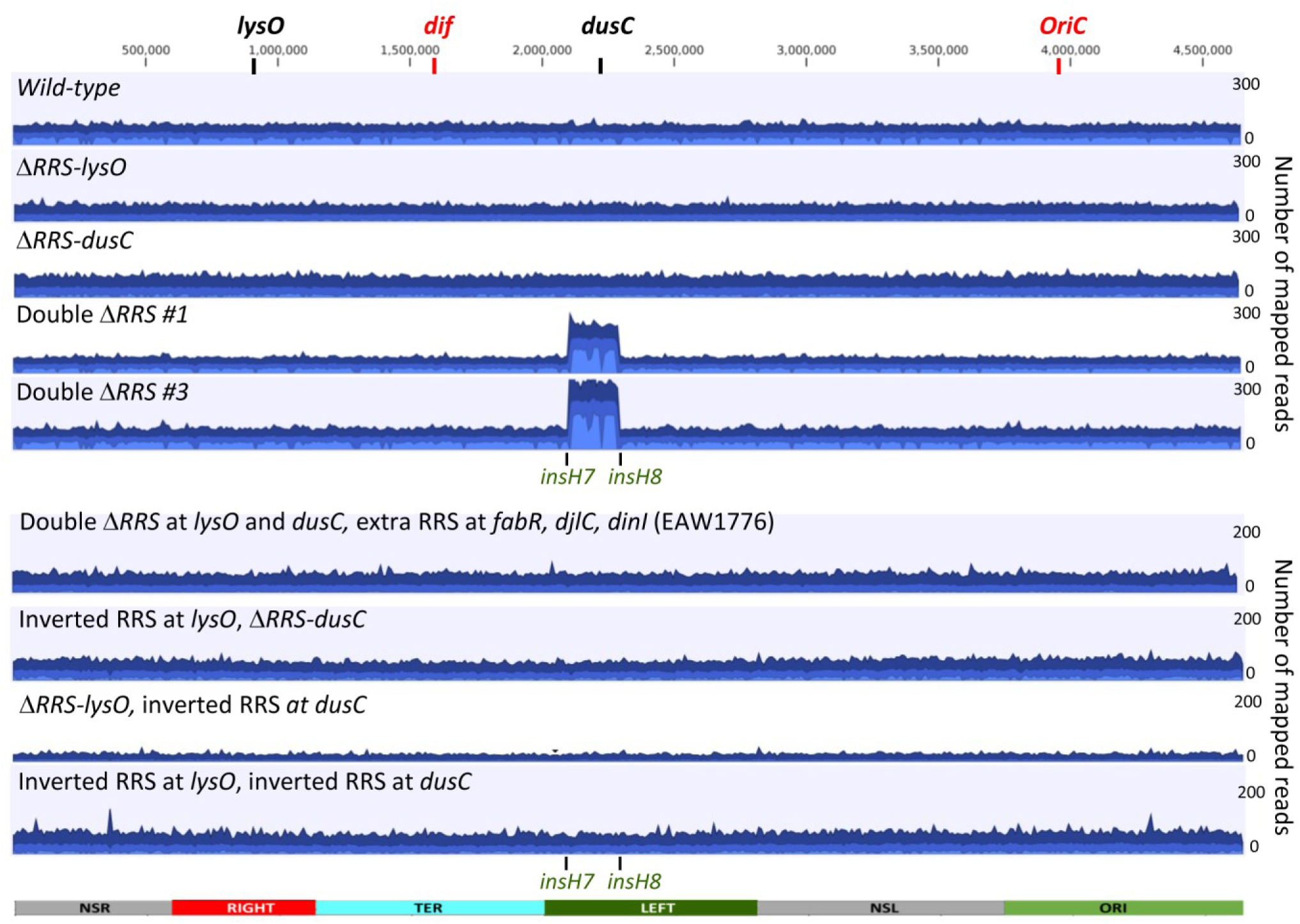
Amplification of *insH7-insH8* genomic region in colonies with double *DRRS-lysO* and *DRRS-dusC* deletion. Whole genome sequencing read mappings of wild-type MG1655, single RRS deletion (*DRRS-lysO* and *DRRS-dusC*) and double *DRRS* deletion strains, and strains with inverted RRS. Surviving double *DRRS* deletion colonies have 2 - 3 folds amplification of a 200 kb region between *insH7* and *insH8* (genome positions from 2102912 to 2288976), encompassing *dusC* locus. Mapped reads are shown as aggregated coverage graphs with average coverage values at each genome position shown in blue, maximum coverage - dark blue and minimum coverage – light blue. Positions of *lys*O, *dusC*, OriC, *dif* and macrodomains are shown on top. Strain EAW1776 contains three RRS elements at *fabR, djlC* and *dinI* loci, but two original RRS at *lysO* and *dusC* had been removed.

We also took EAW1768 strain with three extra RRS at *fabR, djlC* and *dinI* and deleted both of the natural RRS sequences within it (strain EAW1776). In this case, both RRS could be deleted without amplification of the *insH7-8* region (Figure 2B and 4). Similarly, strains containing at least one RRS in the reverse orientation also did not show the *insH7-8* amplification (Figure 4). This result indicates that one or more of the three abnormally positioned RRS elements, or an inverted RRS can compensate for the absence of the natural RRS sites.

The functional complementation of the RRS deletions appeared to be complete. The strains with both RRS deleted, compensated by either the *insH7-8* amplification or the addition of additional RRS, exhibited no growth defect (Figure 5A). In addition, none of the strains exhibited increased sensitivity to either UV irradiation or ciprofloxacin (Figure 5B).

**Figure 5.**
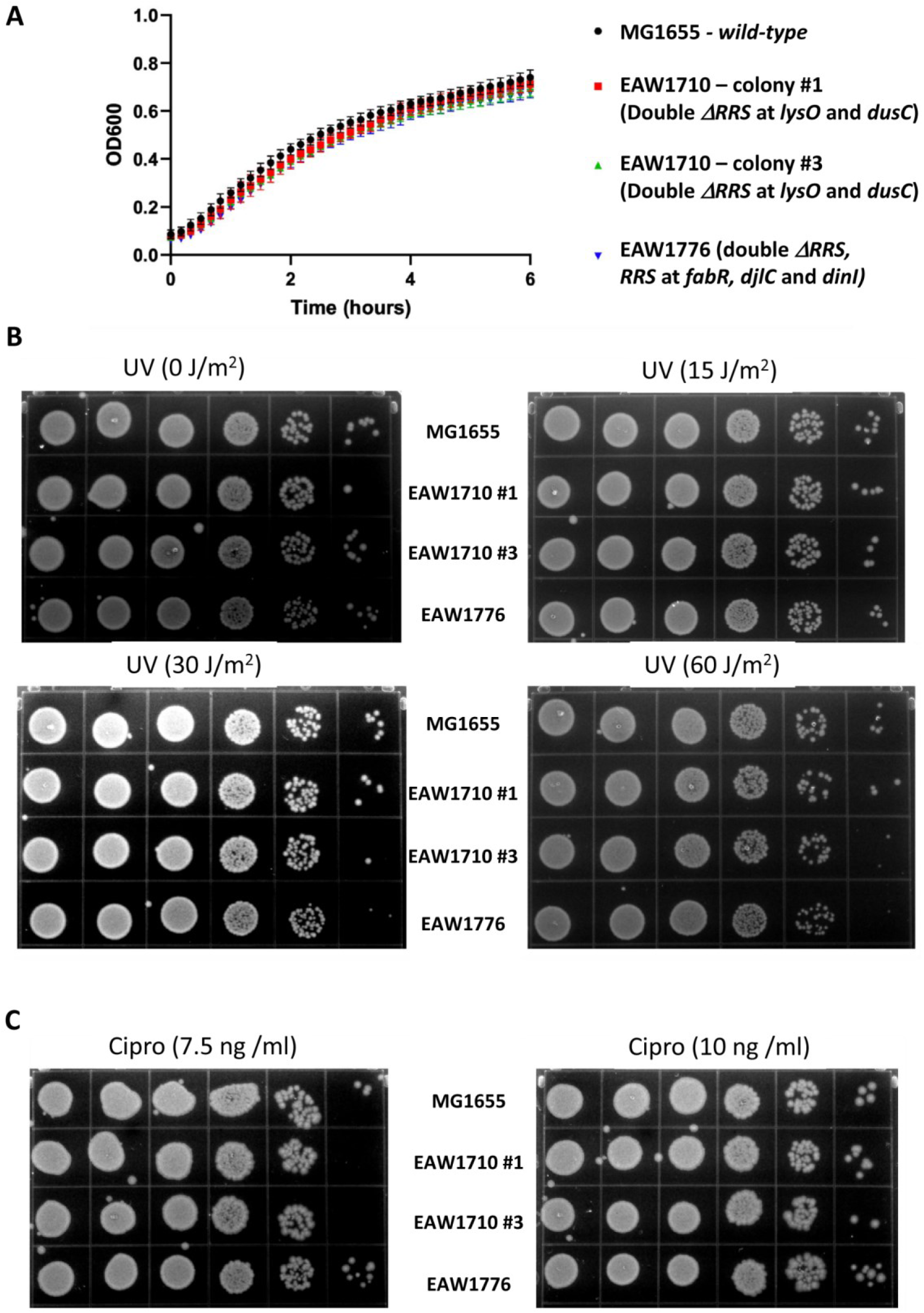
Deletion of both RRS (with compensation) does not affect growth or sensitivity to UV or ciprofloxacin. **A**, Growth curves for wild-type and double *ΔRRS* deletion strains. Two colonies of EAW1710 (double Δ*RRS* deletion), each picked separately, are strains with the *insH7-8* region amplified but otherwise have no detectable RRS elements. EAW1776 has both of the natural RRS deleted but also has three additional RRS in other genomic locations as shown in Figure 2A. Panels in parts **B** and **C** show UV and ciprofloxacin sensitivities, respectively.

### RRS elements affect genome-wide supercoiling

To examine effects of RRS deletions on chromosome supercoiling genome-wide, we employed a new method developed by Bates and coworkers, called Psora-seq (24). The method measures relative supercoiling density (σ) by crosslinking of biotin-tagged psoralen to the genomic DNA *in vivo*, and then carrying out deep sequencing to determine the extent of psoralen binding along the chromosome. Psoralen binds preferentially to negatively supercoiled DNA (33,34), and the rate of binding is proportional to negative superhelical density of DNA (35,36). Psora-seq allows identification of genomic regions that are more negatively supercoiled (psoralen-enriched regions with high binding z-scores) or are more positively supercoiled (psoralen-depleted regions with low z-scores) in comparison to the genome σ average (Figure 6A). Overall, the circular *E. coli* chromosome is slightly negatively supercoiled with an estimated average σ = −0.05 (36).

**Figure 6.**
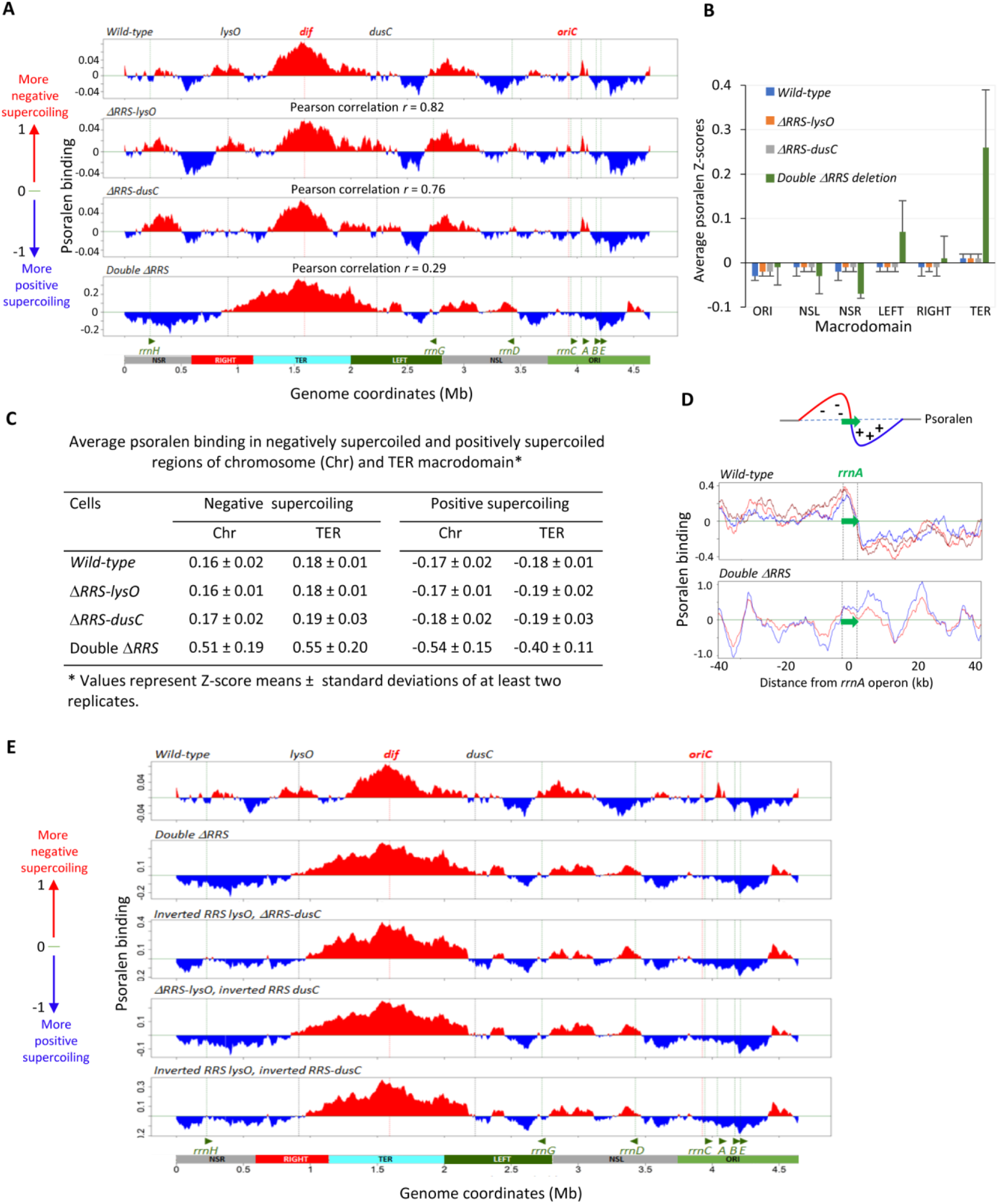
RRS elements affect genome-wide supercoiling. **A,** Psora-seq maps (250 kb resolution) of *wild-type* and RRS deletion mutants. Psoralen binding values (log2 pull-down/input) are the average of at least two independent Psora-seq experiments for mid-exponential cells. Red and blue tracks indicate supercoiling that is more negative or more positive than the genome average, respectively. Pearson correlation coefficients (*r*) are shown for comparison between psoralen binding profiles of RRS deletion mutants and *wild-type*. Positions OriC, *dif, lysO, dusC* and 7 ribosomal operons (*rrn*) are indicated by vertical dashed lines. **B**, Quantification of psoralen binding within macrodomains of wild-type and RRS deletion mutants. Error bars indicate ± standard deviations. **C**, Average psoralen binding in negatively and supercoiled regions of whole chromosome and TER macrodomain. Psoralen binding z scores in **B** and **C** were calculated using 1 kb sliding windows. **D**, The absence of the twin supercoiling domain signature at *rrn* operons in double *DRRS* cells. Psoralen binding along an 80 kbp region surrounding *rrnA* operon from three independent replicates for wild-type (*top graph*) and two replicates for double *DRRS* cells (*bottom graph*). Sketch on top showed predicted psoralen binding profile according to a transcription-induced supercoiling twin-domain model. **E**, Comparison of large-scale supercoiling features of *E. coli* chromosomes (250 kb resolution) from wild-type, double D*RRS* deletion, single D*RRS* deletion with an inverted *RRS* and double inverted *RRS* strains.

When applied to the *E. coli* genome (Figure 6), the published results (24) were closely replicated with wild type cells. In particular, there is a region in the TER macrodomain where increased negative supercoiling is observed. Similar patterns of genomic supercoiling were observed when either one of the two native RRS were deleted. However, when both RRS are deleted (compensated for by the *insH7-8* amplification), major changes in global genome topology are observed. The effect is seen most dramatically when results are averaged over 250 kb windows. The region near the terminus in which negative supercoiling is substantially increased is expanded, now covering the entire region extending between the sites of the deleted RRS (*lysO-dusC*) (Figure 6A). Additional effects are seen throughout the genome. When results are averaged over smaller windows (Supplementary Figures S5 and S6), more details become evident. Psoralen binding profiles of single RRS deletion strains are highly correlated (*r* ∼ 0.8) with the wild-type profile, whereas double Δ*RRS* deleted cells have a much lower correlation value (*r* = 0.3) (Figure 6A). Average psoralen binding z-scores across macrodomains vary from - 0.03 to 0.02 in wild-type and singe RRS deleted cells (Figure 6B). However, a much higher z-score variation (from −0.07 to 0.26) between macrodomains was observed in double Δ*RRS* cells with notable enrichment of bound psoralen in TER macrodomain (Figures 6B). Significantly higher levels of supercoiling in double Δ*RRS* cells are also evident when psoralen binding z-scores were calculated separately for negative and positive supercoiling genomic regions (Figure 6C).

The *E. coli* genome supercoiling landscape is largely determined by transcription with an observed distinctive twin-supercoiling domain signature at ribosomal (*rrn*) operons (24). Twin-supercoiling domains are formed when an elongating RNA polymerase causes overwinding of downstream DNA and underwinding of upstream DNA, generating positively and negatively supercoiled regions, respectively (37) (Figure 6D, *top sketch*). While the twin-domain signature was present at all seven *rrn* operons for wild-type and single RRS deletion strains, it was not observed in double Δ*RRS* deleted cells (Figure 6D, Supplementary Figure S7). The absence of the *rrn* twin supercoiling pattern suggests that there may be additional mechanisms, such as those involving DNA replication (38,39), which may exert a more dominant role in defining genome supercoiling in double Δ*RRS* cells.

The strains with RRS elements inverted were also subjected to Psora-seq (Figure 6E). In this case, the results closely mirrored the results seen when both RRS were deleted and complemented by the *insH7-8* genome amplification. The region near the replication terminus again featured large increases in negative supercoiling extending for the entire 1.3 Mb distance between the normal locations of the native RRS.

DNA in the *E. coli* genome is organized into multiple independent supercoiled domains such that different levels of superhelical density σ can be maintained in different regions of the same chromosome (40,41). These topological domains may result from the presence of topological insulators that impede supercoiling diffusion into different domains (42,43).

Although the identities and positions of topological insulators along the chromosome have yet to be established, bacterial interspersed mosaic elements such as type 2 (BIME-2) have been implicated as topological barriers. BIME-2 elements are in the intergenic regions and serve as targets for DNA gyrase *in vivo* (14). Insertion of a BIME-2 element into the chromosome has been shown to prevent the propagation of transcription induced supercoiling along a reporter sequence (43). In *wild-type* MG1655 there are 208 BIME-2 sites (14), which do not distribute uniformly along the chromosome, with only few sites present in the TER macrodomain (Figure 7A). Examination of our Psora-seq data around BIME-2 sites revealed that most BIME-2 sites (70% - 77%) are located in between supercoiling regions, compared to 23% to 30% in supercoiling regions (Figure 7A and 7B). The high percentage of BIME-2 observed in the border between supercoiling genomic regions, even in double *ΔRRS* cells (Figure 7A and 7B), indicate that these elements seem likely to serve as *bone fide* topological barriers that impede DNA supercoiling propagation occurring not only during transcription, but also during DNA replication.

**Figure 7.**
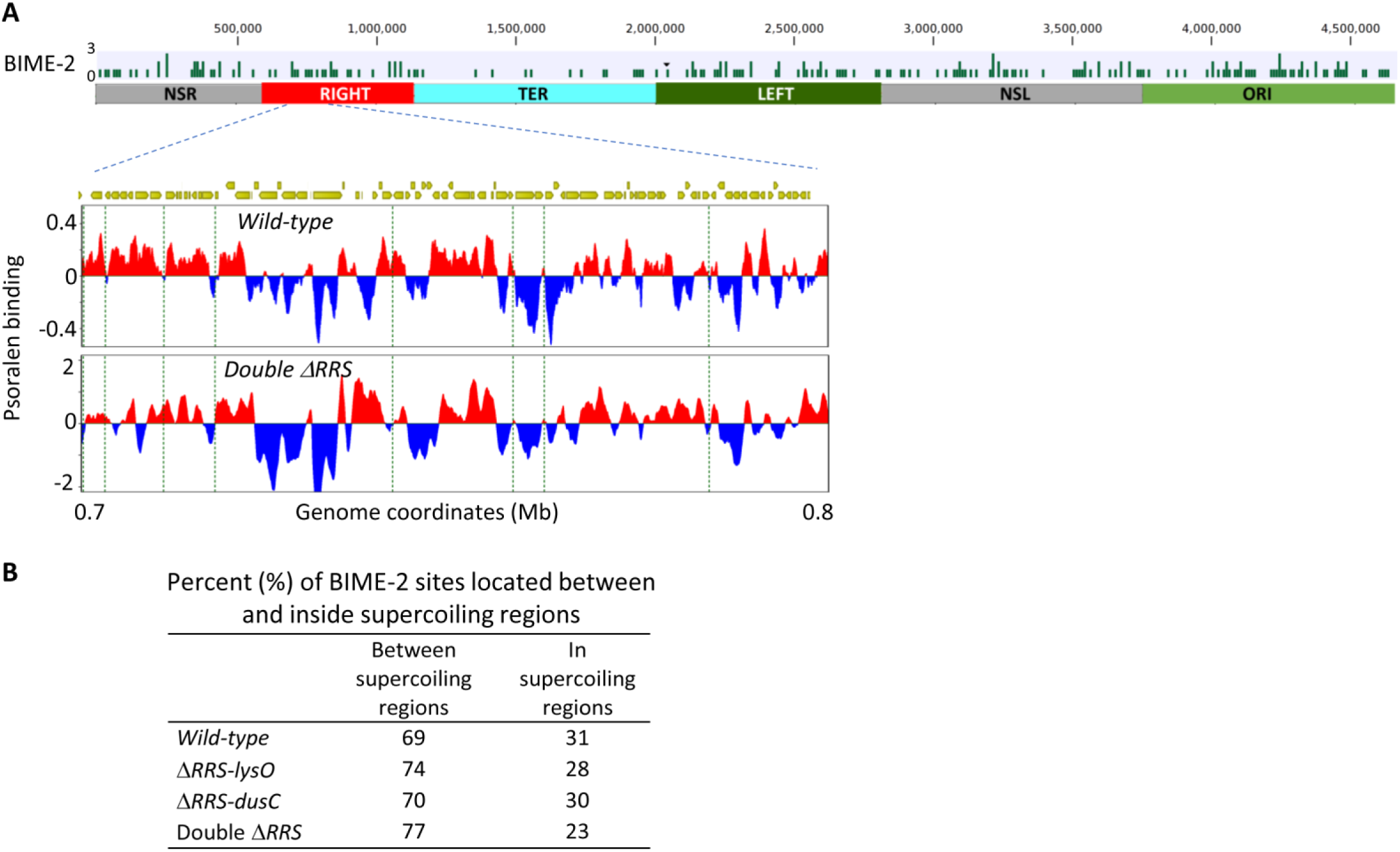
Most BIME-2 sites are located between supercoiling regions of *E. coli* chromosome. **A**, Distribution of BIME-2 sites along *E. coli* chromosome. Each bar in the graphs represents the number of BIME-2 sites in a 10 kb genomic segment (*top graph*). Psora-seq maps in genomic region between 0.7 - 0.8 Mb for wild-type and double *ΔRRS* (*bottom graphs*). Positions of BIME-2 sites are indicated by green vertical dashed lines. **B**, Percentage of BIME-2 sites located between and inside supercoiling regions.

## DISCUSSION

The RRS is a genomic sequence that triggers gap formation on the lagging strand (Figure 5), a gap into which SSB and RecA are often loaded. This study establishes that (a) the RRS has a substantial effect on replication, particularly when the G4-containing strand is used as template, (b) the RRS functions at any genomic location in which it is placed, (c) at least one RRS is conditionally essential (both can be deleted at their native locations only if a large genomic segment is amplified or they are replaced by inverted RRS elements), (d) RRS at least some other locations may also substitute for the RRS at their normal genomic location, and (e) removal or inversion of both RRS has a dramatic effect on overall genomic topology.

The function of the RRS element at other genomic locations is telegraphed by the deposition of RecA protein at the new locations as seen with ssGap-seq. We have not been able to cleanly delete both RRS and obtain a viable cell. Such deletions are always accompanied by genomic amplification of a large region encompassing one of the RRS. Determining how this amplification can compensate for the missing RRS function will require additional research.

RRS at other genomic locations can compensate for deletion of both natural RRS elements. In our experiments, three additional RRS were present in the genome. Two of these are somewhat close to the natural RRS present near *lysO*. More work will be needed to determine if any one of these RRS is sufficient on its own to compensate for the nature RRS elements. Finally, both RRS can be replaced by inverted versions of the RRS. However, deletion of both RRS (with the *insH7-8* region amplification), as well as replacement of the RRS sequences with inverted RRS, has a dramatic effect on global genome topology. These results speak to a likely role for the RRS elements in genome topology maintenance.

The effects of deleting both RRS on genome topology is easy to document. We hypothesize that normal RRS function is related to genome topological transitions during replication termination and chromosome segregation at cell division. Formation of a transient gap would have the effect of introducing potential swivels at every phosphodiester bond in the ssDNA region. These could relax DNA stress related to supercoiling or they might facilitate some aspect of chromosome folding. Additional support for the proposed role of transient gaps in regulation of genome topology came from Psora-seq data for strains with inverted RRS elements. The absence of ssDNA gaps at *lysO* and *dusC* loci in the inverted RRS strains (Figure 3, Supplementary Figure S2) caused changes in the chromosome supercoiling landscape, which resemble the changes observed for the double Δ*RRS* deletion strain (Figure 6E).

We assume that gap formation is transient, as conversion of the gap to duplex DNA would eventually be necessary for subsequent replication cycles. The formation of a transient gap would introduce potential swivels at every phosphodiester bond that could relieve topological stress. We postulate that the RRS provide topological relief valves, facilitating the complex genomic transitions that accompany replication termination and chromosome segregation. The RRS could also be the structural manifestation of the TER macrodomain boundary, a function that would not exclude additional roles.

Global patterns of genome topology are largely transcription-driven in wild type cells (24). This pattern is disrupted when both RRS elements are removed from the genome. This observation in no way undermines the concept of transcription-driven genome topology in the wild-type genome. However, it does suggest that the RRS elements have some undetermined role in maintaining this regime. There may be layers of control over nucleoid topology. The RRS elements may provide topological relief to an extent in which the effects of transcription, particularly at *rrn* operons, are dominant and readily observed. Those same transcription effects may simply be somewhat masked when overall topology is constrained by the removal of both RRS elements. In double Δ*RRS* cells, we speculate that there could be additional mechanisms, perhaps involving DNA replication (38,39), that exert a predominant role in supercoiling.

Bulky DNA lesions routinely trigger the formation of lesion-containing post-replication gaps, which are generally repaired either via RecF-mediated recombinational DNA repair or translesion DNA synthesis (44). The presence of genomic elements that trigger post-replication gap formation has not previously been considered, primarily because nobody has looked for them. One other example of such an element has appeared in the bacterial literature, a G-quadruplex-forming structure that triggers gap formation and is required for pilin antigenic variation in *Neisseria gonorrhoeae* (45,46). In principle, any DNA sequence that can fold up quickly – a G-quadruplex or any other stable tertiary structure – as duplex DNA is unwound during replication, could potentially block full replication of the lagging strand template and create a transient gap (Supplementary Figure S8). Sequences with the potential to form G-quadruplexes are common in eukaryotic genomes, with approximately 700,000 such sequences in the human genome (47–52). Triggering transient gap formation, in the service of chromosome structural transitions or replication, could be a function of some or many of them, perhaps to facilitate chromosome folding or relieve topological stress.

## DATA AVAILABILITY

Raw next-generation sequencing data have been deposited to SRA database (SRA Submission ID: SUB13887136, BioProject ID: PRJNA1025028). BioProject’s metadata is available at: https://urldefense.com/v3/_https://dataview.ncbi.nlm.nih.gov/object/PRJNA1025028?reviewer=77hfshm2vmjkc71kbmim7dbcvv;!!LIr3w8kk_Xxm!pNUteqXxvIvwIkexT_5kA5o9yPS0y5fs8vRs2PjHADuovPNcxfywVmqg4wXZ8nMKSGcRinJKeF0iOs7I$

## SUPPLEMENTARY DATA

Supplementary Data are available at NAR Online.

## ACKNOWLEDGMENTS

This work was supported by grants from National Institutes of General Medical Sciences (1RM1GM130450 & 1R21AI179102 to M.M.C., M.F.G.) and National Institute of Environmental Health Sciences (R35ES028343 to M.F.G.). We thank Joseph Purwanto for his assistance in preparation of *E. coli* cells for ssGap-seq and Psora-seq experiments.

