## Supplementary Figure S1 for "Controlling Genome Topology with Sequences that Trigger Post-replication Gap Formation During Replisome Passage: The *E. coli* RRS Elements"

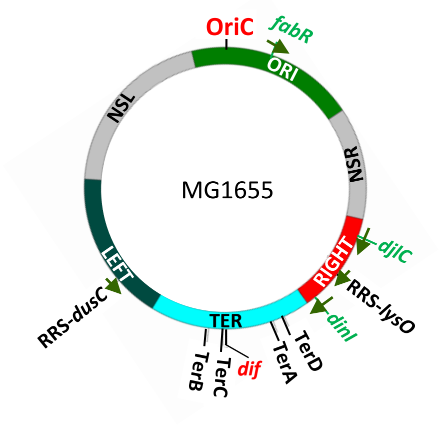


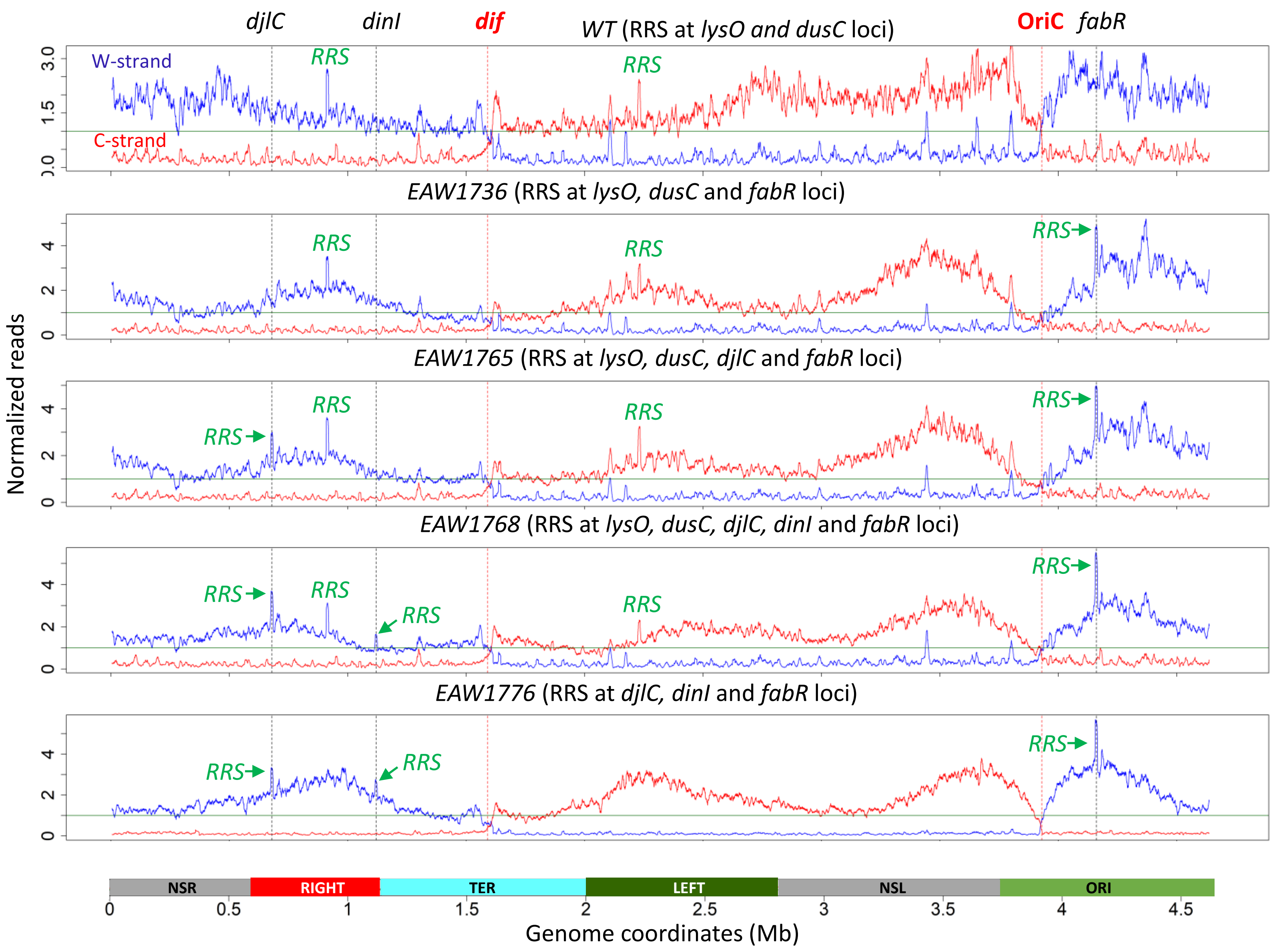


**Supplementary Figure S1. Genome-wide distribution of SSB-ssDNA in *wild-type* and mutant strains containing extra RRS sequences inserted at *fabR* locus (EAW1736); *fabR* and *djlC* loci (EAW1765); *fabR, djlC* and *dinI* loci (EAW1768).** Strain EAW1776 contains three RRS elements at *fabR, djlC* and *dinI* loci, but two original RRS at *lysO* and *dusC* had been removed. Normalized coverages of SSB-ssDNA are shown as blue traces for the W-strand, and red traces for the C-strand. Green lines in graphs (value = 1) indicate the genome average coverage. Each data point represents a moving average of SSB-ssDNA in a 10 kb window.
