## Supplementary Figure S2 for "Controlling Genome Topology with Sequences that Trigger Post-replication Gap Formation During Replisome Passage: The *E. coli* RRS Elements"

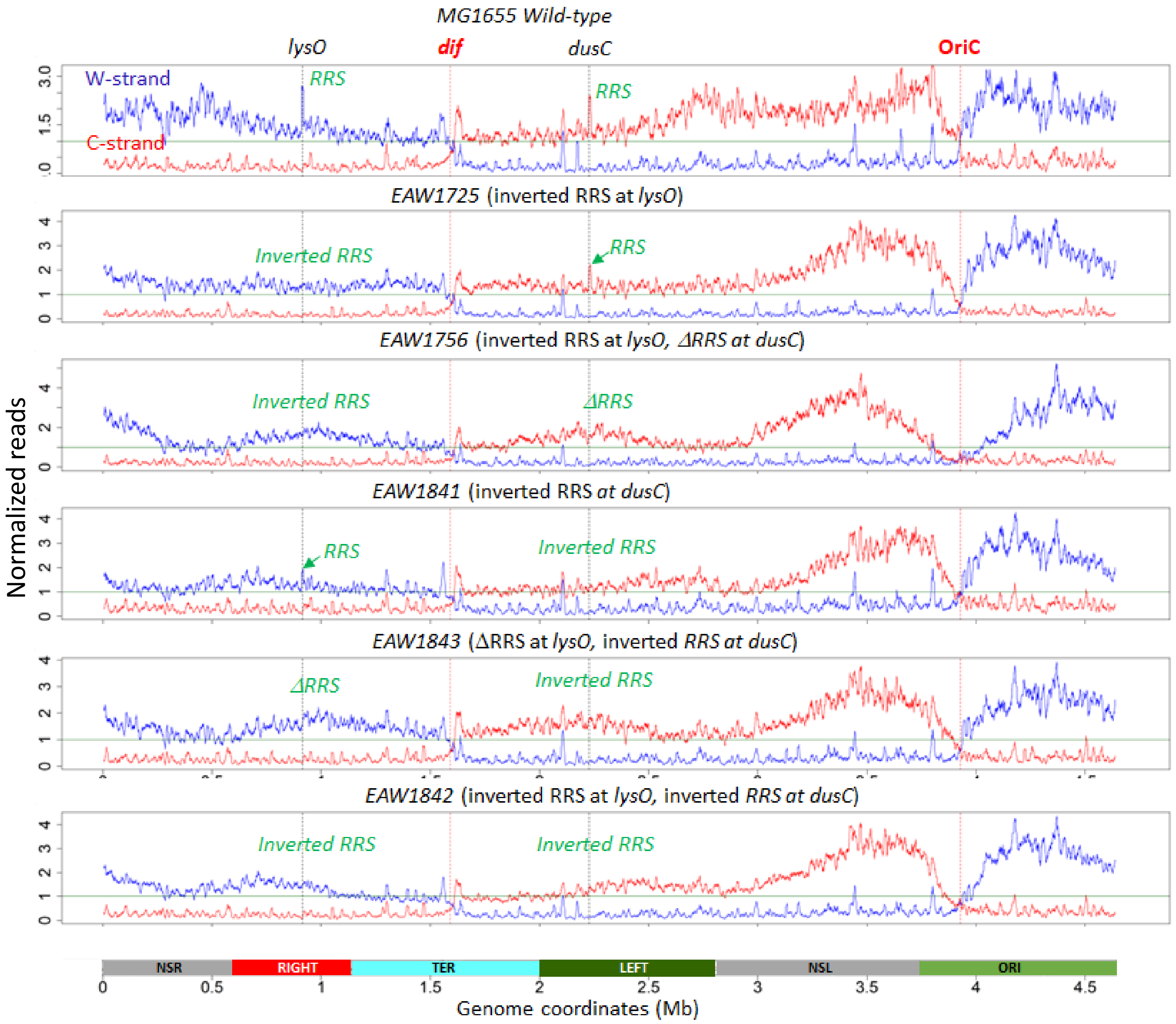


**Supplementary Figure S2. Genome-wide distribution of SSB-ssDNA in *wild-type* and mutant strains with inverted RRS: an inverted *RRS-lysO* (EAW1725), an inverted *RRS-lysO* and ΔRRS deletion at *dusC* (EAW1756), an inverted *RRS-dusC* (EAW1841), an inverted *RRS-dusC* and Δ*RRS* deletion at *lysO* (EAW1843*)*, and double inverted *RRS-lysO* and *RRS-dusC* (EAW1842).** Normalized coverages of SSB-ssDNA are shown as blue traces for the W-strand, and red traces for the C-strand. Green lines in graphs (value = 1) indicate the genome average coverage. Each data point represents a moving average of SSB-ssDNA in a 10 kb window.
