## Supplementary Figure S3 for "Controlling Genome Topology with Sequences that Trigger Post-replication Gap Formation During Replisome Passage: The *E. coli* RRS Elements"

**A**

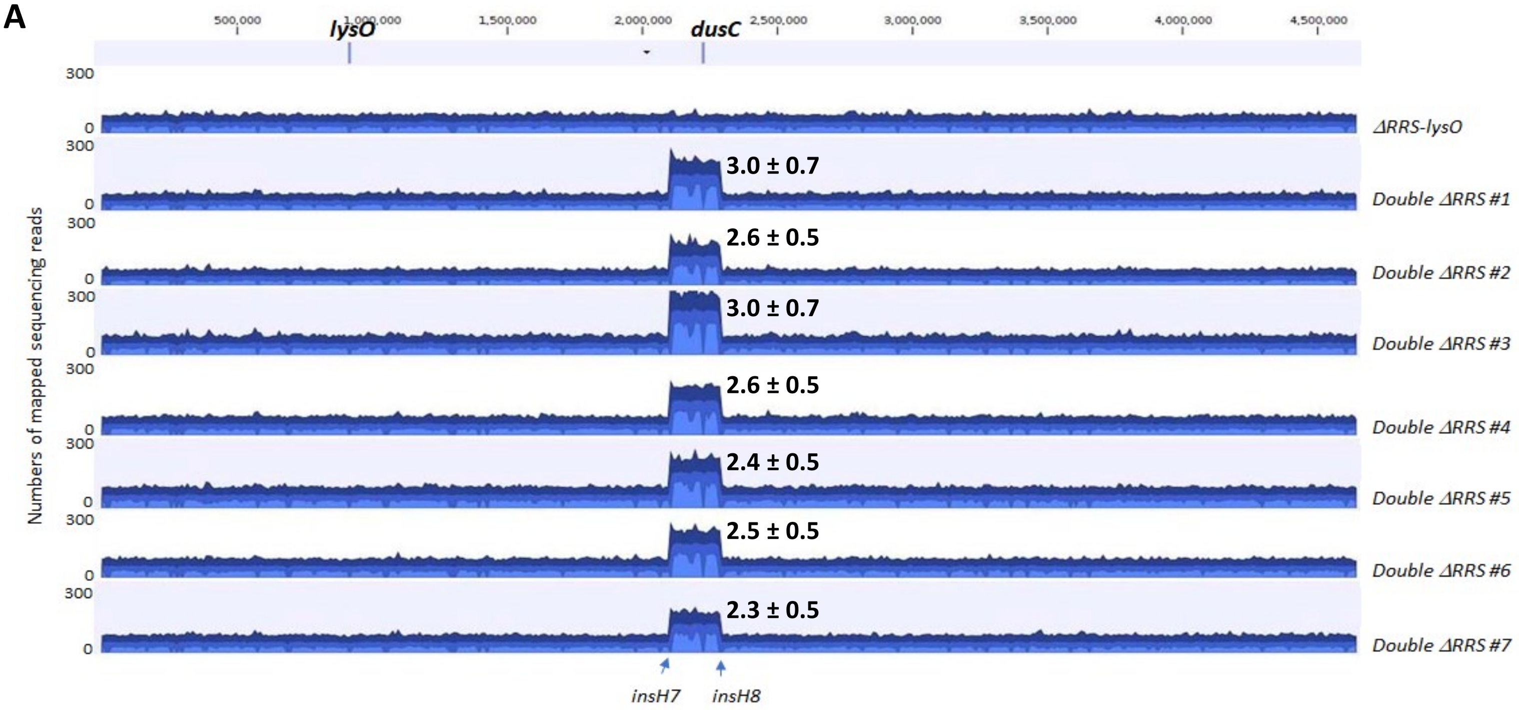


**B**

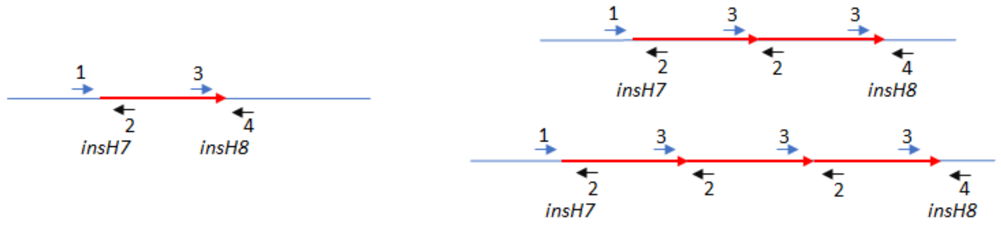


**Single Δ*RRS* deletion strains EAW1706 & EAW1708**

Expected PCR products:
Primer 1 + primer 2 (lanes **L**): Yes
Primer 3 + primer 4 (lanes **R**): Yes
Primer 2 + primer 3 (lanes **T**): No

**Four double Δ*RRS* deletion colonies EAW1710**

Expected PCR products:
Primer 1 + primer 2 (lanes **L**): Yes
Primer 3 + primer 4 (lanes **R**): Yes
Primer 2 + primer 3 (lanes **T**): Yes

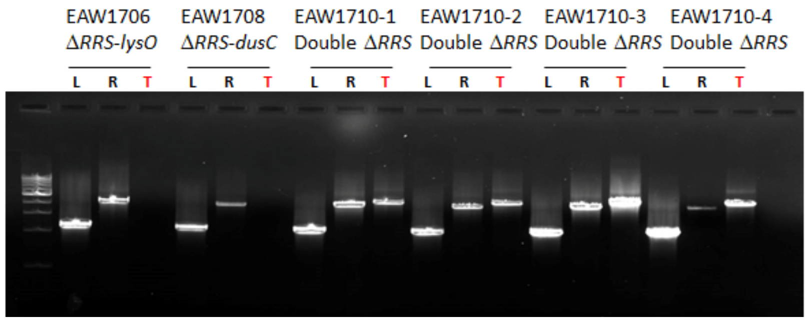


**Supplementary Figure S3. Tandem duplication/triplication of *insH7-insH8* genomic region in colonies with double Δ*RRS-lysO* and Δ*RRS-dusC* deletion. A,** Whole genome sequencing read mappings of a single RRS deletion (ΔRRS-*lysO*) and seven double Δ*RRS* colonies (Double Δ*RRS* #1 to #7). Surviving double Δ*RRS* deletion colonies have 2 - 3 folds amplification of a 200 kb region between *insH7* and *insH8*, encompassing *dusC* locus. Mapped reads are shown as aggregated coverage graphs with average coverage values at each genome position shown in blue, maximum coverage - dark blue and minimum coverage – light blue. The Y-axis gives the numbers of mapped reads. The locations of *lysO* and *dusC* loci are indicated on the top. The numbers on the top of the *insH7-8* region represent the average numbers of sequencing reads in the region compared to the genome average (means ± standard deviations). **B**, PCR product patterns are consistent with the presence of tandem duplication/triplication of the *insH7-insH8* genomic region in four double Δ*RRS* colonies.
