## Supplementary Figure S4 for "Controlling Genome Topology with Sequences that Trigger Post-replication Gap Formation During Replisome Passage: The *E. coli* RRS Elements"

**A**

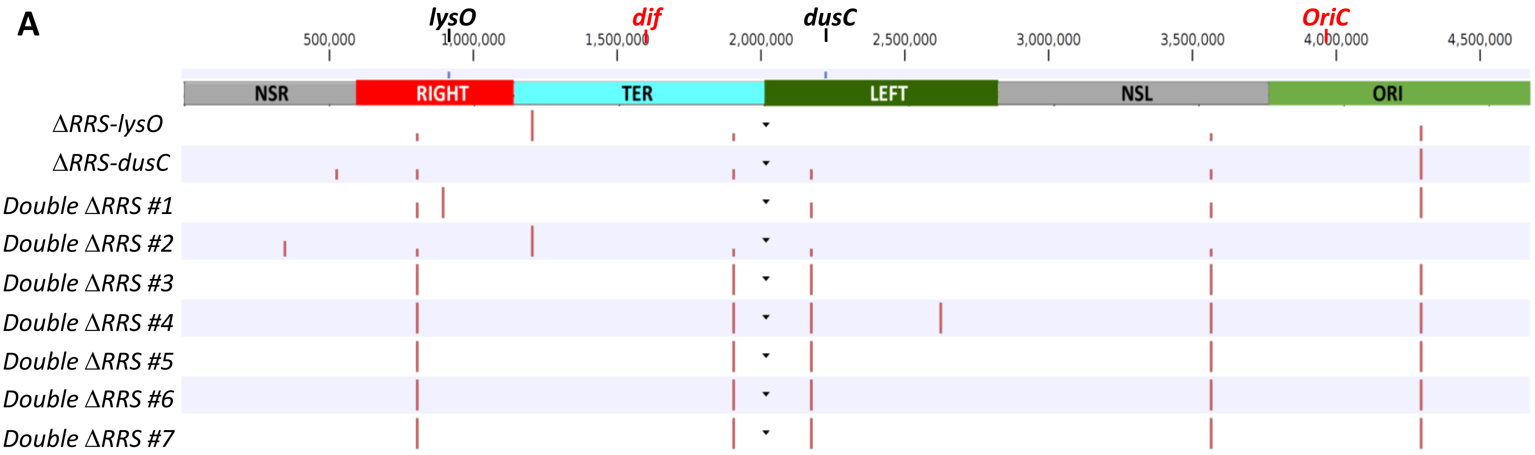


**B**

| Position | | *Single ΔRRS deletion* | | | | Double Δ*RRS* deletion | | | | |
|---|---|---|---|---|---|---|---|---|---|---|
| | Mutation (*gene*, AA change) | *lysO* | *dusC* | 1 | 2 | 3 | 4 | 5 | 6 | 7 |
| 343796 | C to T (*yahK,* D305Y) | - | - | - | + | - | - | - | - | - |
| 524036 | C deletion (*rhsD*) | - | + | - | - | - | - | - | - | - |
| 803662 | C to A (*ybhJ,* L54I) | + | + | + | + | + | + | + | + | + |
| 896791 | G to T (*potH,* D220Y) | - | - | + | - | - | - | - | - | - |
| 1207789 | C to G (*ycfK,* L97V) | + | - | - | + | - | - | - | - | - |
| 1209619 | C to G (*intergenic*) | + | - | - | + | - | - | - | - | - |
| 1905761 | G to A (*mntP,* G25D) | + | + | - | + | + | + | + | + | + |
| 2173361 | CC deletion (*integenic*) | - | + | + | + | + | + | + | + | + |
| 2621437 | G to A (*purM,* D81N) | - | - | - | - | - | + | - | - | - |
| 3560455 | G insertion (*intergenic*) | + | + | + | + | + | + | + | + | + |
| 4296060 | C to T *(intergenic)* | + | + | + | - | - | - | - | - | - |
| 4296380 | CG insertion (*intergenic*) | - | + | - | - | + | + | + | + | + |

**Supplementary Figure S4. Single nucleotide variant (SNV) analysis for MG1655 mutant colonies with a single and double Δ*RRS* deletions. A,** Genomic locations of detected SNV variants are shown as vertical red lines. **B**, Annotated list of detected SNV in the sequenced colonies. SNV variants were identified using the “Identify DNA Germline Variants” workflow in CLC Genomic workbench (v. 23.0.1).
