## Supplementary Figure S5 for "Controlling Genome Topology with Sequences that Trigger Post-replication Gap Formation During Replisome Passage: The *E. coli* RRS Elements"

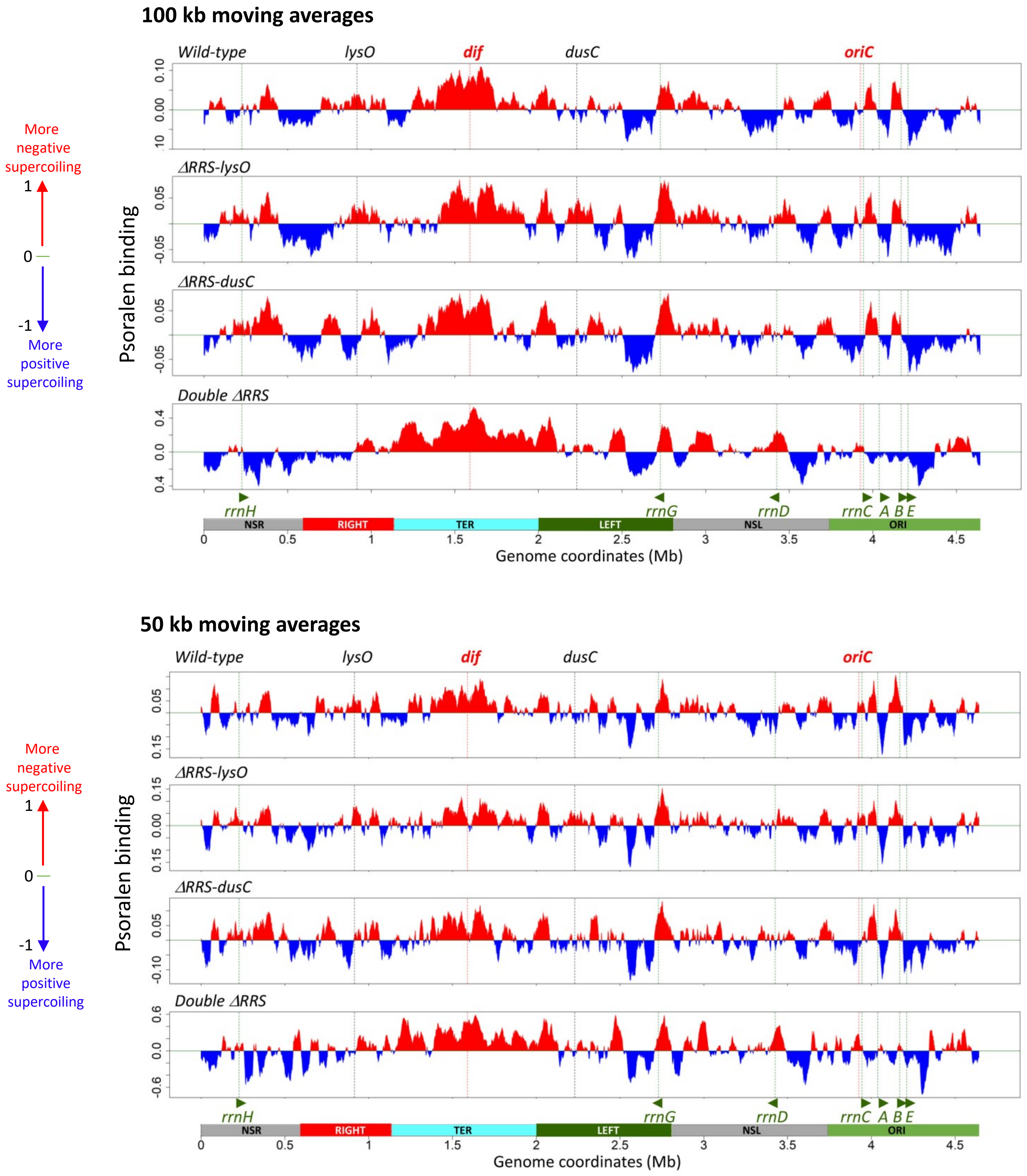


**Supplementary Figure S5. Large-scale supercoiling features of *E. coli* chromosomes from wild-type, single and double Δ*RRS* deletion strains in mid-exponential growth phase**. Psora-seq maps at 100 kb and 50 kb resolutions. Psoralen binding values were calculated as log2 (pull-down/input). Red and blue tracks indicate supercoiling that is more negative or more positive than the genome average, respectively. Positions of *oriC*, *dif, lysO, dusC* loci are indicated on the top, *rrn* operons and macrodomains are shown at the bottom.
