## Supplementary Figure S7 for "Controlling Genome Topology with Sequences that Trigger Post-replication Gap Formation During Replisome Passage: The *E. coli* RRS Elements"

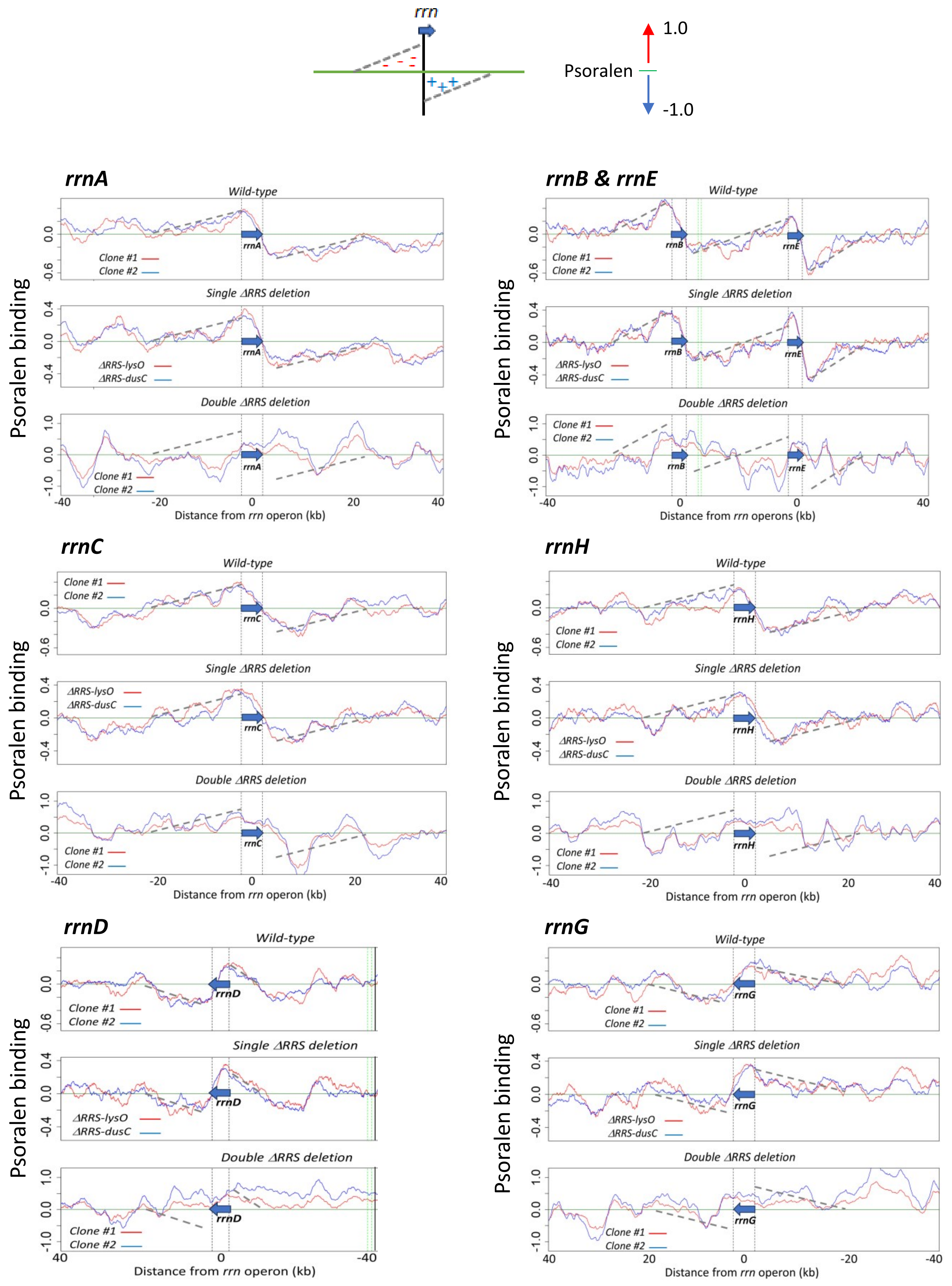


**Supplementary Figure S7. The absence of twin-supercoiling domains at *rrn* operons in double *ΔRRS* cells.** Psoralen binding to 40 kb regions upstream and downstream of ribosomal RNA operons *rrnA*-*G*. Twin-domains of supercoiling are formed when an elongating RNA polymerase causes overwinding of downstream DNA and underwinding of upstream DNA, generating positively and negatively supercoiled regions, respectively (*top sketch*). Psoralen binding patterns in 25 kb regions around *rrn* operons are consistent with the presence of twin-supercoiling domains in wild-type and single Δ*RRS* deleted cell, but not in double Δ*RRS* cells. Positions of *rrn* operons are indicated by vertical black dashed lines.
