## Supplementary figures and images for "Controlling Genome Topology with Sequences that Trigger Post-replication Gap Formation During Replisome Passage: The *E. coli* RRS Elements"

### Supplementary Figure S8

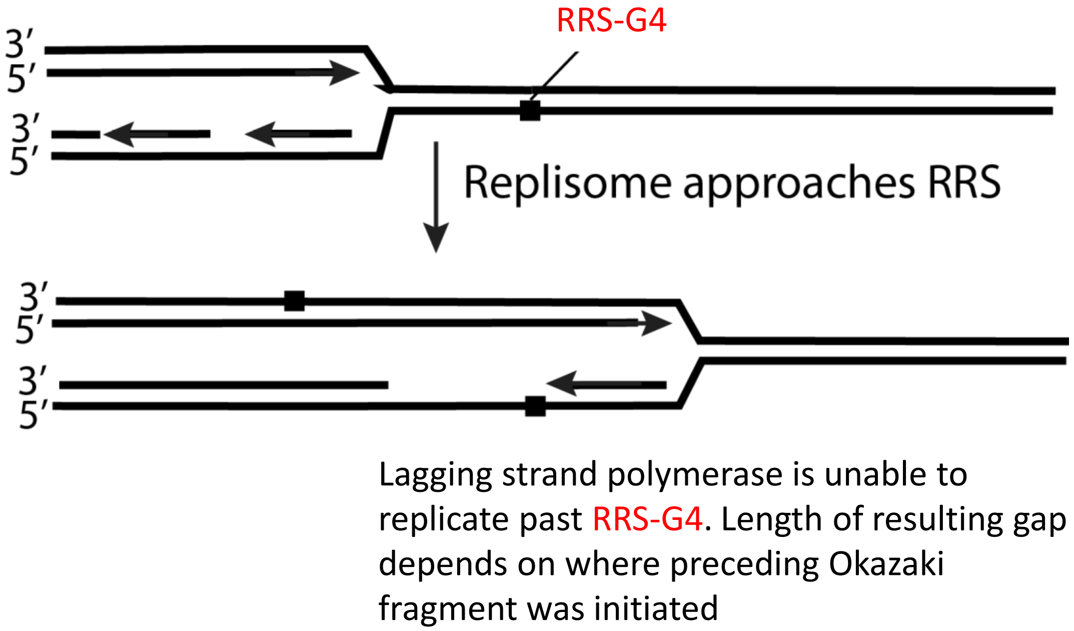


**Supplementary Figure S8. Model for how RRS triggers gap formation on the lagging strand.**
